# Boymaw, Overexpressed in Brains with Major Psychiatric Disorders, May Encode a Small Protein to Inhibit Mitochondrial Function and Protein Translation

**DOI:** 10.1101/005728

**Authors:** Baohu Ji, Minjung Kim, Kerin K. Higa, Xianjin Zhou

## Abstract

The t(1,11) chromosome translocation co-segregates with major psychiatric disorders in a large Scottish family. The translocation disrupts the DISC1 and Boymaw (DISC1FP1) genes on chromosomes 1 and 11, respectively. After translocation, two fusion genes are generated. Our recent studies found that the DISC1-Boymaw fusion protein is localized in mitochondria and inhibits oxidoreductase activity, rRNA expression, and protein translation. Mice carrying the DISC1-Boymaw fusion genes display intermediate behavioral phenotypes related to major psychiatric disorders. Here, we report that the Boymaw gene encodes a small protein predominantly localized in mitochondria. The Boymaw protein inhibits oxidoreductase activity, rRNA expression, and protein translation in the same way as the DISC1-Boymaw fusion protein. Interestingly, Boymaw expression is up-regulated by different stressors at RNA and/or protein translational levels. In addition, we found that Boymaw RNA expression is significantly increased in the postmortem brains of patients with major psychiatric disorders. Our studies therefore suggest that the Boymaw gene is a potential susceptibility gene for major psychiatric disorders in both the Scottish t(1,11) family and the general population of patients.

## Introduction

Major psychiatric disorders are severe brain disorders which cause a heavy burden to both society and family. Due to the absence of biological hallmarks, it is difficult to investigate the underlying molecular mechanisms and to diagnose and treat these disorders in clinics. Classical genetic studies, however, identified a high penetrance genetic mutation from a large Scottish family in which the t(1,11) chromosome translocation co-segregates with major psychiatric disorders across five generations [1,2]. There has been extensive interest in studying this family for various reasons. First, identification of a high penetrance susceptibility gene from the family may provide key insights into our understanding of the underlying molecular mechanisms of psychiatric disorders. Second, such a susceptibility gene may also contribute to the development of psychiatric disorders in the general population. Over the last decade, disruption of the DISC1 gene by the chromosome translocation has been the focus of the studies on the Scottish family [3]. However, our previous studies found that the translocation disrupts not only the DISC1 gene, but also the Boymaw (DISC1FP1) gene, and consequently results in the formation of the DISC1-Boymaw and Boymaw-DISC1 fusion genes [4–6]. Recently, we found that the DISC1-Boymaw fusion protein reduces oxidoreductase activities, rRNA expression, and protein translation in both *in vitro* cells and in mice carrying DISC1-Boymaw fusion genes (manuscript submitted). Surprisingly, fusion of the Boymaw gene to a randomly selected gene generates the same cellular phenotypes as the DISC1-Boymaw fusion gene, suggesting that the Boymaw gene is likely responsible for the inhibition of oxidoreductase activity and of protein translation. These findings prompted us to investigate the roles of the Boymaw gene.

In the present study, we report that the Boymaw gene encodes a small protein predominantly localized in mitochondria. We examined the functions of the Boymaw gene and its expression in response to stress. To explore whether Boymaw could be a potential susceptibility gene for major psychiatric disorders, we quantified Boymaw RNA expression in the postmortem brains of patients with schizophrenia, bipolar disorder, and major depression from the general population.

## Methods and Materials

### DNA constructs

All Boymaw expression DNA constructs are driven by a CMV promoter and terminated with an SV40 poly(A) signal in a pTimer-1 plasmid vector (Clontech) containing a fluorescence timer (FT) gene. The Boymaw-HA plasmid was constructed by a fusion of an HA tag at the C-terminus of the full-length Boymaw ORF3. The Boymaw-HA-mut plasmid was generated by a mutation of the ATG start codon of uORF2 in the Boymaw-HA plasmid. The Boymaw-HA-ACG plasmid was created by a mutation of the ATG start codon of ORF3 in the Boymaw-HA plasmid. An HA epitope and fused FT was inserted 10 amino acids downstream of the ATG start codon of the Boymaw ORF3 to replace the rest of the Boymaw ORF3 in the Boymaw-HA-FT. Boymaw-HA-FT-mut construct was generated by a mutation of the ATG start codon of uORF2. All DNA constructs were completely sequenced.

### *In vitro* cell culture

HEK293T (ATCC, CRL-11268) and COS-7 cells (ATCC, CRL-1651) were cultured in DMEM or DMEM/F12 medium supplemented with 10% fetal bovine serum, penicillin, and streptomycin (Life Technologies, CA) at 37°C in a humidified atmosphere containing 5% CO_2_. HEK293T and COS-7 cell transfections were performed using Mirus TransIT® reagent (Mirus, WI), according to manufacturer’s instructions.

### ITRAQ analysis

After starvation, HEK293T cells were harvested and sent to UCSD Mass Spectrometry Facility Core for iTRAQ analysis. To replicate the experiments, the cells were split into two samples which were labeled with 114 Da and 115 Da, mass tags respectively. Protein identification and quantification were performed using the Paragon™ Algorithm in ProteinPilot™ Software. Putative Boymaw ORF3 protein sequence was included into the database for searching.

### *MTT reduction* assays

MTT (3-(4, 5-dimethylthiazol-2-yl)-2, 5-diphenyl tetrazolium bromide reduction assays were conducted as described [7] with slight modifications. In brief, HEK293T cells were seeded at cell density of 1 × 10^5^ cells per well in 24-well plates, 24 h prior to transfection. Cell proliferation was measured by counting cell numbers with 0.04% Trypan blue 48 h after transfection. MTT was added to the cell lysate at the final concentration of 0.5 mg/ml, and the reaction was incubated at 37 °C for 1 h. MTT crystals were collected and dissolved in DMSO. Absorbance at 540 nm was measured with SpectraMax M5 Multi-Mode Microplate Readers (Molecular Devices, PA).

### Western blot and Immunocytochemical analyses

Western blot analyses were conducted as described [8]. In brief, total proteins were extracted from HEK293T cells in Passive Lysis Buffer (Promega, WI) containing 0.2% Sarkosyl and 1X protease inhibitor cocktail (Sigma, MO) at room temperature for 15 min on a shaker. Protein concentration was measured with Bradford (Abs 595 nm) method using Coomassie Plus Protein Assay (Thermo Scientific, IL). After electrophoresis, proteins were transferred onto PVDF membranes. Rat monoclonal anti-HA antibody (1:500) conjugated with peroxidase (3F10) (Roche, CA) was used to directly detect HA epitopes. The same membrane was stripped and re-probed with mouse monoclonal anti-β-actin (1:5,000), sc-47778 (Santa Cruz Biotechnology, CA). After washing, the membranes were further incubated with horseradish peroxidase-conjugated anti-mouse IgG (1:5,000, Cell signaling, MA) for 1.5 h at room temperature.

Quantification of protein expression was conducted with Image J. For immunocytochemical staining, COS-7 cells were transfected with TransIT®-COS Transfection reagent and COS Boss Reagent. 40 h after transfection, cells were fixed with 4% paraformaldehyde/PBS at room temperature for 10 min. The fixed cells were blocked with 2% goat serum/PBS containing 0.2% Triton-X100 at room temperature for 1 h, and further incubated with 1:250 dilution of rabbit anti-HA antibody (H6908, Sigma-Aldrich) overnight. After rinsing with PBST three times, the cells were incubated with Alexa Fluor®488 goat anti-rabbit IgG (1:500, Invitrogen) for 1 h at room temperature. After washing with PBST, the cells were mounted with VECTASHIELD® Hard Set™ Mounting Medium containing DAPI (Vector, CA). For analysis of subcellular localization of the Boymaw ORF3 protein in COS-7 cells, the following primary antibodies were used: mouse monoclonal anti-Cytochrome C (1:250), ab110325 (Abcam, MA); rabbit polyclonal anti-HA (1:400), H-6908 (Sigma-Aldrich, MO). The secondary antibodies are: Alexa Fluor^®^ 568 goat anti-mouse IgG (1:500), (Invitrogen, CA) and Alexa Fluor^®^ 488 goat anti-rabbit IgG (1:500), (Invitrogen, CA). The coverslips were mounted with VECTASHIELDR Hard Set™ Mounting Medium with DAPI (Vector, CA). Images were acquired with an Olympus FluoView™ FV1000 confocal microscope.

### SUnSET

The SUnSET experiments were conducted as described [9]. In brief, HEK293 cells seeded at cell density of 1 × 10^5^ cells per well in 24-well plates, 24 h prior to transfection. After transfection with the Boymaw-HA, Boymaw-HA-ACG, and Boymaw-HA-mut plasmids, cells were pulse-labeled with puromycin at the concentration of 10 ug/ml for 10 min. Total proteins were extracted and loaded on PAGE gels (10ug per lane) for Western blot analysis. Anti-puromycin antibody (1:10,000)(12D10, Millipore) was used to detect incorporation of puromycin in newly synthesized proteins. Blots were stripped later and re-probed with ant-β-actin antibody. Image J was used to quantify the incorporation of puromycin.

### Starvation

HEK293T cells were seeded in 12-well plates 20 h prior to transfection. 1.2 µg of each plasmid DNA and 3.6 µL of TransIT®-293 Transfection were mixed to transfect cells in each well. 16 h after transfection, the culture medium was replaced with either fresh culture medium for the controls or Earle’s Balanced Salt Solution for amino acids starvation.

### Hypoxia

HEK293T cells were seeded in 12-well plates 20 h prior to transfection. 16 h after transfection, hypoxia was induced by the addition of CoCl_2_ at a final concentration of 100 uM. Total proteins were extracted at 0, 16, 24, and 32 h after CoCl2 treatment. Western blot was conducted as described.

### Real-Time PCR

In all real-time PCR experiments, we confirmed the specific amplification of PCR DNA bands on agarose gel. All amplified DNA bands were specific and confirmed by sequencing. Total RNA was extracted from cells with TRIzol reagent (Invitrogen, Carlsbad, CA). cDNA was synthesized from 5ug of total RNA using Superscript III First-Strand Synthesis System (Invitrogen, Carlsbad, CA) with random hexamers. SYBR Green real-time PCR was used to quantify relative expression of the Boymaw ORF3 and HIF-1α RNA with a comparative Ct method. β-actin was used as a reference control. The standard curve of PCR amplification was first examined. All amplifications had R^2^ >0.99. Amplification efficiency for each pair of primers was determined using known molecular concentration of template DNA. Each sample had 4 amplification replica wells. After amplification, *Ct* was determined in the exponential amplification phase of the curve as recommended by the manufacturer’s protocol (Bio-Rad CFX384). Variation of cycles between amplification replica wells was smaller than 0.5 cycles. To quantify the Boymaw ORF3 RNA in human brains, real-time PCR was used against two different reference controls. In the first quantification experiment, one-step real-time PCR using Power SYBR® Green RNA-to-CT™ 1-Step Kit (Invitrogen, Carlsbad, CA) was used with 18S rRNA as a reference control. In the second quantification experiment, two-step real-time PCR was employed. 1.2 µg total RNA was first converted into cDNA using Superscript III First-Strand Synthesis System (Invitrogen) with random hexamers. SYBR Green was used to quantify relative expression of the Boymaw ORF3 RNA with a comparative Ct method. β-actin was used as a reference control. The following primers were used in real-time PCR:

18SRNA-F: CGGCTTTGGTGACTCTAGATAAC; 18SRNA-R: GTGGACTCATTCCAATTACAGG; Boymaw-F: AGCTTAAGACGTACTCCTCAAGGGG; Boymaw-R: CCTAAGACCCACAGATGGAAT; HIF-1α-F: TCCATGTGACCATGAGGAAA; HIF-1α-R: CCAAGCAGGTCATAGGTGGT; β-actin-F: TTCTACAATGAGCTGCGTGTG; β-actin-R: GGGGTGTTGAAGGTCTCAAA; DISC1-F: GCAGCCAGCTCTTAGCAGTT; DISC1-R: CTCCATTCTCAGTGGGGTGT.

### RNA from postmortem human brains

We received 57 BA7 cortical RNA samples out of 60 brains from Stanley Medical Research Institute (SMRI), consisting of 15 each diagnosed with schizophrenia, bipolar disorder, or major depression, and 15 unaffected controls. The four groups were matched by age, sex, race, postmortem interval, pH, hemisphere, and mRNA quality (http://www.stanleyresearch.org/dnn/Default.aspx?tabid=196).

### Statistical analysis

All data were first tested for normal distribution using the Kolmogorov–Smirnov test. ANOVA was used for statistical analysis of all data from cell culture studies. For statistical analyses of the Boymaw RNA expression in postmortem human brains, ANOVA with diagnostic code as a between-subjects factor was performed. *Post hoc* analyses were carried out using Newman-Keuls or Tukey’s test. Alpha level was set to 0.05. All statistical analyses were carried out using BMDP statistical software (Statistical Solutions Inc., Saugus, MA).

## Results

The Boymaw gene is evolutionarily conserved in exon-intron splicing junctions (GT-AG) in primates (Supplemental Figure 1a). Exons 3 to 6 of the human Boymaw gene contain the largest Open Reading Frame 3 (ORF3) with the potential to encode 63 amino acid residues (red). In the Scottish family, the translocation breakpoint is localized in intron 3. After translocation, 59 out of 63 amino acid residues of the ORF3 are in-frame fused at the C-terminus of the truncated DISC1 to form the DISC1-Boymaw fusion gene [4]. Although the fused Boymaw ORF3 can be translated in the form of the DISC1-Boymaw fusion protein, it is unknown whether endogenous Boymaw ORF3 naturally produces any protein. In addition to the ORF3 (red), there are two smaller upstream ORFs (uORF1 and uORF2, blue) in human Boymaw RNA (Figure 1a). After aligning the Boymaw gene in primates, we found that the translational start codon (ATG) is conserved only in ORF3, but not in uORF1 and uORF2. Additionally, high homology is found between the putative proteins encoded by the ORF3 in primates (Supplemental Figure 1b). The structural organization of the Boymaw gene indicates that the ORF3 of the Boymaw gene may encode a small protein.

**Figure 1.**
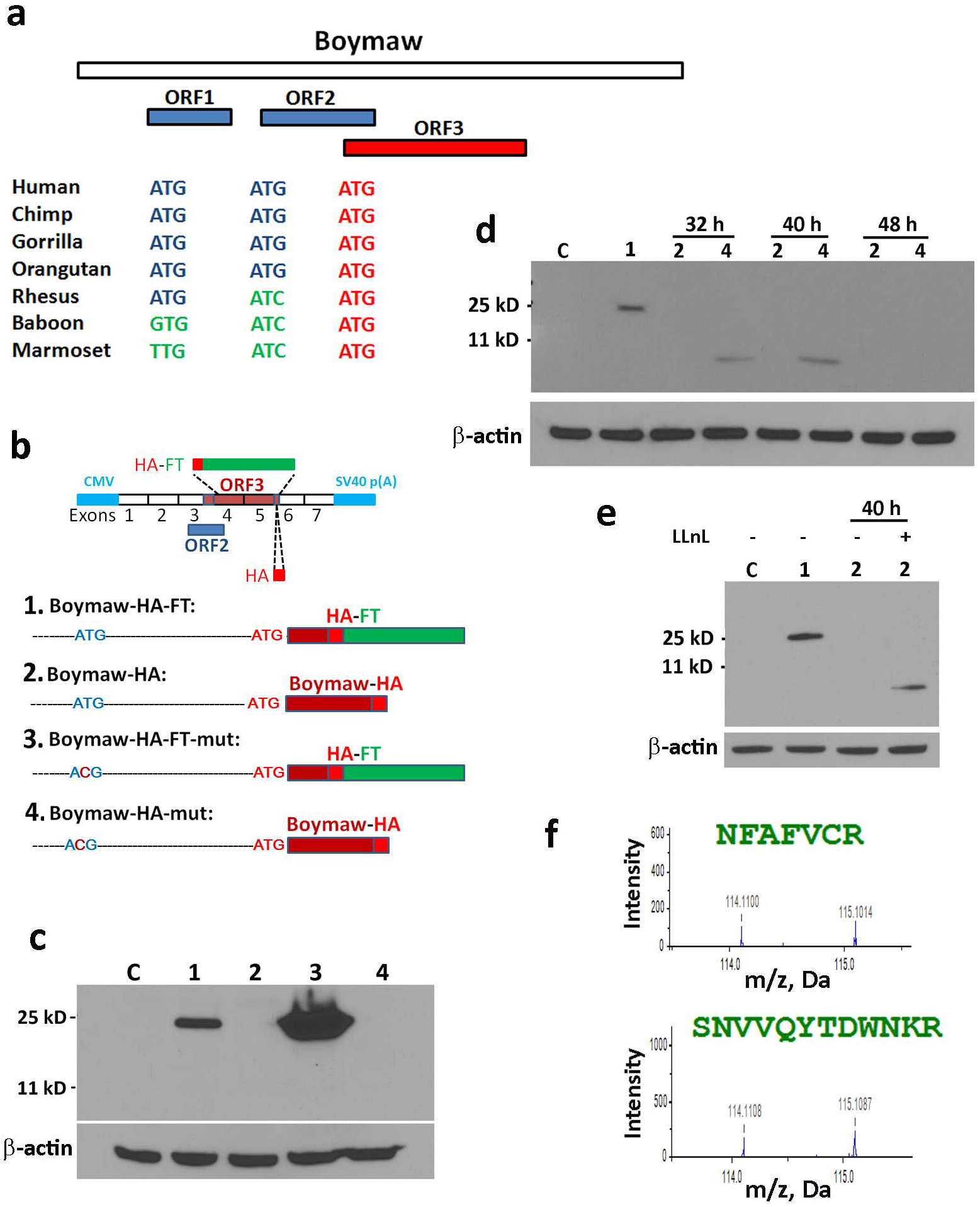
Boymaw ORF3 encodes a small protein. **(a)** There are two ORFs (uORF1 and the uORF2) upstream of the largest ORF3 in the Boymaw RNA. Only the ATG start codon of the ORF3, not the uORF1 and the uORF2, are conserved in primates. **(b)** The four DNA constructs (1, 2, 3, 4) are driven by a CMV promoter and an SV40 poly(A) in a pTimer plasmid vector. In both the Boymaw-HA-FT and Boymaw-HA-FT-mut plasmids, HA epitope tag and fluorescence timer (FT) protein are in-frame fused 10 amino acids downstream of the ATG start codon of ORF3. In the Boymaw-HA and the Boymaw-HA-mut plasmids, HA epitope tag is in-frame fused at the C-terminus of ORF3. In both the Boymaw-HA-FT-mut and the Boymaw-HA-mut plasmids, the ATG start codon of the uORF2 is mutated into ACG. **(c)** Western blot analyses of the HA tagged proteins from the four plasmid constructs. Protein expression was readily detected, 48 h post-transfection, from cells transfected with either the Boymaw-HA-FT or the Boymaw-HA-FT-mut constructs, but not from the Boymaw-HA and the Boymaw-HA-mut constructs. Equal amounts of proteins were loaded (see β-actin expression). C: control cells without plasmid transfection; 1, 2, 3, 4: cells transfected with each of the four different plasmids. **(d)** The HA tagged full-length Boymaw ORF3 protein (about 7 kD) was detected from the cells transfected with the Boymaw-HA-mut plasmids at 32 h and 40 h (but not 48 h post-transfection), suggesting rapid degradation of the full-length Boymaw ORF3 proteins. Equal amounts of proteins were loaded (see β-actin expression). C: control cells without plasmid transfection; 1: positive control cells transfected with the Boymaw-HA-FT plasmids; 2 and 4: cells transfected with Boymaw-HA and Boymaw-HA-mut plasmid, respectively. **(e)** Proteasome inhibitor N-acetyl-L-leucyl-L-leucyl-L-norleucinal (LLnL) was added to inhibit protein degradation at the final concentration of 40 uM to increase the level of the full-length Boymaw ORF3 proteins in cells transfected with the Boymaw-HA construct. Western blot detected the full-length Boymaw ORF3 proteins 40 h post-transfection in the presence of the LLnL. C: control cells without plasmid transfection; 1: positive control cells transfected with the Boymaw-HA-FT plasmids; 2: cells transfected with the Boymaw-HA plasmids. **(f)** Two Boymaw peptides “NFAFVCR” and “SNVVQYTDWNKR” were detected by iTRAQ analysis of human HEK293T cells, suggesting that endogenous Boymaw ORF3 produce a small protein.

## Boymaw ORF3 Encodes a Small Protein

To investigate whether the Boymaw ORF3 is translated in human cells, we in-frame fused an HA-tagged fluorescence timer (FT) protein downstream of the ORF3 start codon in the Boymaw-HA-FT construct (Figure 1b). Translation of the ORF3 would produce an HA-tagged fusion protein with molecular weight of 25 kD (there is no other internal ATG start codon that can generate HA-tagged proteins). However, insertion of a long HA-FT sequence may potentially alter RNA secondary structure surrounding the ORF3 start codon to cause artificial translation. To minimize any potential structural alteration from the insertion of the HA-FT sequence, we fused a short HA epitope tag (27 nucleotides) at the C-terminus of the ORF3 to generate the Boymaw-HA construct. It has been demonstrated that upstream ORFs (uORFs) inhibit downstream major ORF translation in many genes [10,11. Particularly, Kozak sequences of Boymaw uORF2 (GCC**ATG**A) and ORF3 (GTC**ATG**T) both display a conservation with the Kozak consensus sequence (A/GNC**ATG**G) surrounding the translational start codon, indicating that uORF2 may potentially inhibit the ORF3 translation. To investigate the possible inhibition, we conducted site-mutagenesis to mutate the uORF2 ATG start codon into ACG in both the Boymaw-HA-FT-mut and the Boymaw-HA-mut constructs. After transfection into HEK293T cells, we detected the expression of the Boymaw-HA-FT fusion protein with an expected size of 25 kD in Western blot analysis (Figure 1c), suggesting that the Boymaw ORF3 is translated in human cells. As expected, mutation of the uORF2 translational start codon remarkably increased the translation of the ORF3 in cells expressing the Boymaw-HA-FT-mut gene. However, we did not detect expression of HA-tagged full-length Boymaw proteins from cells transfected with either the Boymaw-HA or Boymaw-HA-mut constructs. We reasoned that the HA tagged full-length Boymaw ORF3 proteins could be unstable and degraded rapidly similar to the DISC1-Boymaw fusion proteins (manuscript submitted). To test our hypothesis, we conducted Western blot to detect HA-tagged full-length Boymaw proteins at earlier time points. In cells transfected with the Boymaw-HA-mut construct, expression of the full-length Boymaw-HA protein (about 7 kD) was detected at 32 h and 40 h, but not 48 h, post-transfection, suggesting a rapid degradation of the full-length Boymaw ORF3 proteins (Figure 1d). However, we did not detect the full-length Boymaw-HA protein from cells transfected with the Boymaw-HA construct. It is possible that the level of the full-length Boymaw-HA protein is lower in cells transfected with the Boymaw-HA construct than the Boymaw-HA-mut construct because the Boymaw uORF2 inhibits the ORF3 translation in the Boymaw-HA construct. To increase the level of the full-length Boymaw-HA proteins in cells transfected with the Boymaw-HA construct, proteasome inhibitor N-acetyl-L-leucyl-L-leucyl-L-norleucinal (LLnL) was added to inhibit protein degradation [12]. As expected, expression of the full-length Boymaw-HA proteins was detected in cells transfected with the Boymaw-HA construct at 40 h post-transfection in the presence of LLnL (Figure 1e). In addition to rapid protein degradation, the small size (7 kD) of the full-length Boymaw protein likely also contributes to weaker signals in Western blot since the blot membrane does not retain small proteins very effectively. To confirm the translation of the Boymaw ORF3 detected by Western blot analysis, we conducted immunocytochemical staining in transfected COS-7 cells (Supplemental Figure 1c). Expression of the Boymaw-HA, Boymaw-HA-mut, Boymaw-HA-FT, and Boymaw-HA-FT-mut proteins were all detected by anti-HA antibodies. To provide direct evidence that Boymaw ORF3 can be translated from human endogenous Boymaw gene, we conducted iTRAQ analysis of human HEK293T cells. Two Boymaw peptides, although their abundances are low, were detected (Figure 1f).

Our previous studies found that the DISC1-Boymaw fusion protein, in which 59 out of 63 amino acid residues of the Boymaw ORF3 are fused at the C-terminus of the truncated DISC1, is predominantly localized in mitochondria in contrast to the diffusive expression pattern of the truncated DISC1 protein (manuscript submitted). The finding prompted us to examine subcellular localization of the full-length Boymaw ORF3 protein. In COS-7 cells transfected with the Boymaw-HA construct, we found that the full-length Boymaw ORF3 protein is almost completely co-localized with cytochrome c, a mitochondrial marker (Figure 2). This mitochondrial localization is not caused by the fused HA epitope since HA-tagged DISC1 and truncated DISC1 proteins do not display co-localization with cytochrome c (manuscript submitted). A z-stack of 15 consecutive optical sections was captured using a confocal microscope (Supplemental Figure 2a). These data suggested that Boymaw ORF3 encodes a small labile protein predominantly localized in mitochondria. This is consistent with our findings (manuscript submitted) and others [6] on the mitochondrial localization of the DISC1-Boymaw fusion protein.

**Figure 2.**
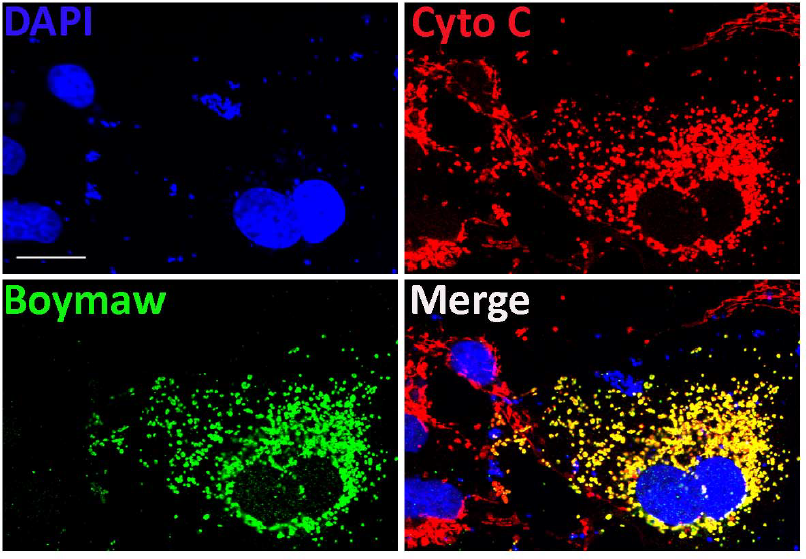
Boymaw ORF3 protein is localized in mitochondria. Co-immunocytochemical stainings of the full-length Boymaw ORF3 protein and cytochrome c. Images were acquired with Olympus FluoView™ FV1000 confocal microscope. Scale bar: 25 µm.

## Boymaw ORF3 Protein Inhibits Protein Translation

Our previous studies demonstrated that the DISC1-Boymaw fusion protein inhibits oxidoreductase activities, rRNA expression, and protein translation (manuscript submitted). We are interested in knowing whether expression of the Boymaw ORF3 protein will generate the same cellular phenotypes as the DISC1-Boymaw fusion protein. To generate a negative control for the studies, we mutated the ORF3 ATG start codon to ACG in the Boymaw-HA-ACG construct to abolish the translation of the ORF3 (Figure 3a). In the Boymaw-HA-mut construct, the uORF2 ATG start codon was mutated to ACG to abolish its inhibition to the downstream ORF3 translation. The Boymaw-HA-ACG and Boymaw-HA-mut constructs differ from the Boymaw-HA in only a single nucleotide. We first compared MTT reduction in cells expressing these genes. As expected, there was no difference in MTT reduction between the constructs when cells were harvested 12 h post-transfection (Figure 3b) because Boymaw ORF3 protein expression requires more than 12 h. However, significant decrease of MTT reduction was observed in cells transfected with either the Boymaw-HA or Boymaw-HA-mut construct at both 40 h and 48 h post-transfection in comparison with either the Boymaw-HA-FT or Boymaw-HA-ACG control.

**Figure 3.**
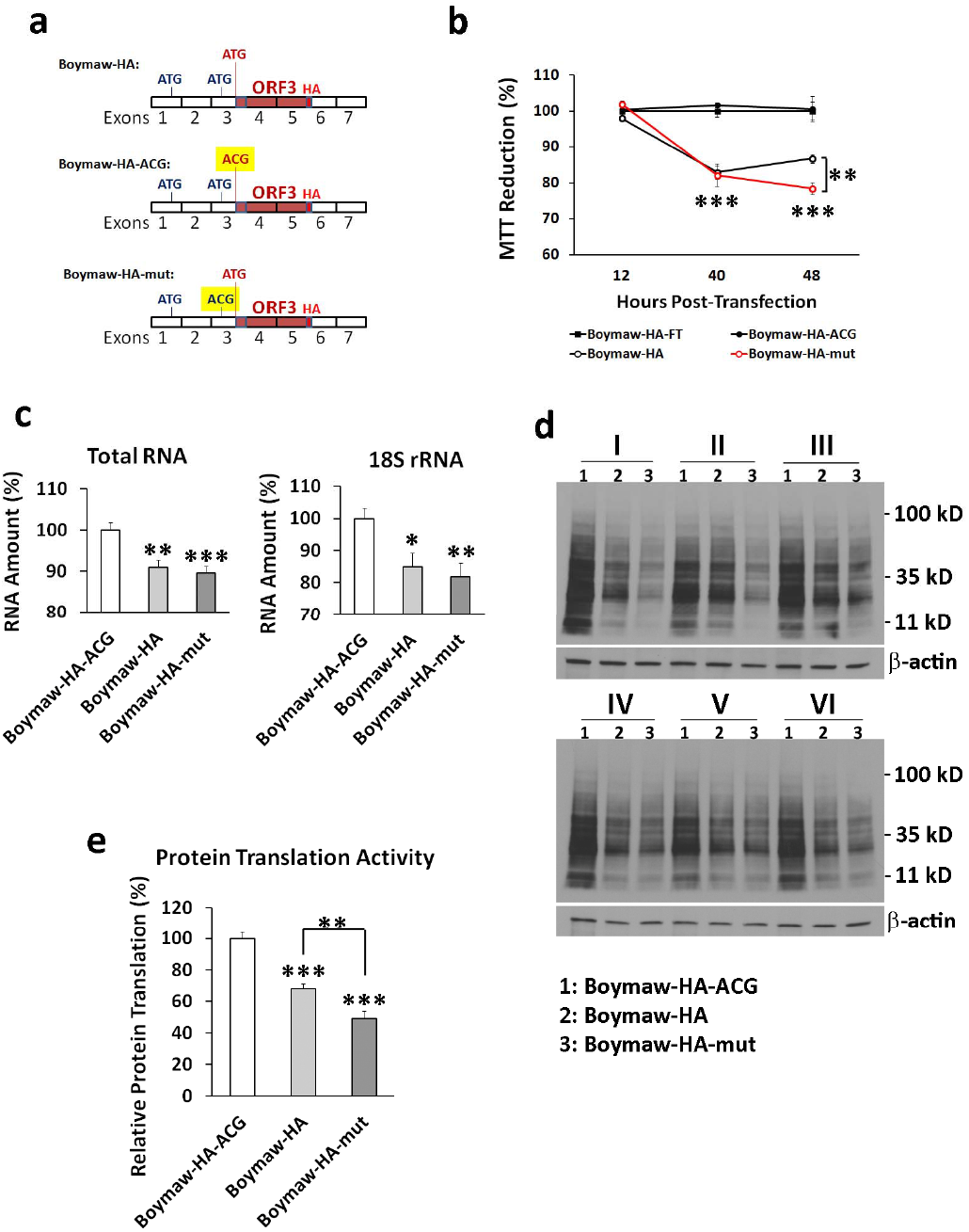
Boymaw ORF3 protein reduces MTT reduction, rRNA expression, and protein synthesis. **(a)** Three Boymaw expression constructs are each driven by a CMV promoter and an SV40 poly(A) in a pTimer plasmid vector. The full-length Boymaw ORF3 protein is tagged with an HA epitope. The ATG start codon of the full-length Boymaw ORF3 was mutated to ACG to abolish the translation of the Boymaw ORF3 in the Boymaw-HA-ACG plasmids. The ATG start codon of the uORF2 was mutated to ACG to abolish its inhibition to the ORF3 translation in the Boymaw-HA-mut plasmids. **(b)** MTT reduction was conducted in cell lysates at different time points after transfection. There were 4 to 5 replica wells per construct per time point. The mean value of MTT reduction in cells transfected with the Boymaw-HA-FT plasmid was used as a control (100%) to calculate relative MTT reduction in the other three constructs. There is a significant gene effect on MTT reduction (F(11,41) = 28. 88, p < 0.00001). *Post hoc* analysis revealed significantly lower MTT reduction in cells transfected with either the Boymaw-HA or the Boymaw-HA-mut constructs at 40 h and 48 h, but not 12 h, post-transfection. There is also significanlyt more MTT reduction in cells transfected with the Boymaw-HA-mut than the Boymaw-HA plasmids at 48 h post-transfection. **(c)** There is a significant gene effect on total RNA (F(2,15) = 11.77, p < 0.001). *Post hoc* analysis revealed a significant reduction of total RNA in cells transfected with either the Boymaw-HA or the Boymaw-HA-mut plasmids in comparison with the Boymaw-HA-ACG control plasmid. There were 6 replica wells per construct. Real-time PCR confirmed a significant gene effect on 18S rRNA expression (F(2,15) = 5.23, p < 0.05). *Post hoc* analysis revealed significant reduction of 18S rRNA in cells transfected with either the Boymaw-HA or the Boymaw-HA-mut. **(d)** SUnSET experiments to measure protein translation activities via puromycin incorporation in cells transfected with the Boymaw-HA, Boymaw-HA-mut, and Boymaw-HA-ACG plasmids. **(e)** Western blot data were quantified with Image J. A significant gene effect on protein translation activities was observed (F(2,15) = 46.85, p < 0.00001). *Post hoc* analysis revealed significant reduction of protein translation in cells transfected with the Boymaw-HA or the Boymaw-HA-mut plasmids in comparison with the control Boymaw-HA-ACG plasmid. Significantly more reduction in protein translation was detected in the Boymaw-HA-mut than in the Boymaw-HA. There were 6 replica wells per construct. Error bar: SEM; * p < 0.05, ** p < 0.01, *** p < 0.001.

Mutation of the ORF3 start codon in the Boymaw-HA-ACG construct generated the same level of MTT reduction as the Boymaw-HA-FT control in which most of the ORF3 is replaced with HA-FT sequence. Therefore, loss of inhibition to MTT reduction by the Boymaw-HA-ACG gene suggests that translation of the ORF3 is necessary for the inhibition. Mutation of the uORF2 start codon in the Boymaw-HA-mut generated more inhibition in MTT reduction than the Boymaw-HA, presumably due to disinhibition of the ORF3 translation. The difference in MTT reduction is not caused by differential cell proliferation (Supplemental Figure 2b). Since MTT reduction is determined by intracellular oxidoreductase activities, the Boymaw ORF3 protein likely inhibits the activities of mitochondrial oxidoreductases to result in decreased MTT reduction. Consistent with the MTT reduction data, transfection of the Boymaw-HA or the Boymaw-HA-mut constructs significantly decreases total RNA expression in comparison with the Boymaw-HA-ACG control (Figure 3c). Real-time PCR quantification confirmed reduction of 18S rRNA expression, likely resulting from overall reduction of rRNA expression. To provide direct evidence for inhibition of protein translation, we conducted SUnSET (surface sensing of translation) experiments to compare protein translation activity in the transfected cells [9]. Significant reduction of protein translation was observed in cells transfected with either the Boymaw-HA or the Boymaw-HA-mut gene in comparison with the Boymaw-HA-ACG control (Figure 3d, 3e). As expected, protein translation was reduced significantly more in cells transfected with the Boymaw-HA-mut construct than the Boymaw-HA construct, presumably due to increased translation of the ORF3 by the uORF2 mutation.

## Induction of Boymaw Expression by Stress

Upstream ORFs (uORF) have been demonstrated to regulate protein translation of major downstream ORFs in response to a variety of stressors such as amino acids starvation and oxidative stress [10]. Therefore, we are interested in investigating a potential regulatory role of the uORF2 in the translation of the Boymaw ORF3. We chose the Boymaw-HA-FT fusion construct instead of the Boymaw-HA construct as a reporter to measure the translation of the Boymaw ORF3, because the full-length Boymaw ORF3 protein generated from the Boymaw-HA construct is unstable and rapidly degraded. The Boymaw-HA-FT-mut construct, in which uORF2 start codon was mutated, was used to investigate the role of uORF2 in the regulation of the ORF3 protein translation. HEK293T cells were transfected with the constructs 16 h before starvation. Translation of the Boymaw ORF3 was examined at different time points post-starvation. Starvation significantly increased the level of the Boymaw-HA-FT fusion protein at 24 h and 32 h post-starvation in comparison with its expression in normal culture medium (Figure 4a). However, mutation of the uORF2 start codon in the Boymaw-HA-FT-mut abolished the increase of the Boymaw-HA-FT fusion protein under starvation. It is possible that alteration of protein turnover may contribute to increased level of the Boymaw-HA-FT protein under starvation. However, we did not observe any increase of the Boymaw-HA-FT protein in cells transfected with the Boymaw-HA-FT-mut construct under starvation, suggesting that protein turnover is not a significant factor for the observed increase of the Boymaw-HA-FT protein expression. These data therefore suggest that increased Boymaw-HA-FT protein expression is caused by increased protein translation of the ORF3 (fused with HA-FT) and that the uORF2 is necessary for the induction of ORF3 translation under starvation. To investigate whether starvation regulates the Boymaw ORF3 expression at the RNA level, we quantified expression of the endogenous Boymaw RNA isoform containing the ORF3 in HEK293T cells. Due to the absence of commercial TaqMan primers to quantify the Boymaw ORF3 RNA expression, we used SYBR Green-based real-time PCR for quantification. The primers used for real-time PCR quantification were first confirmed by amplification and sequencing of the correct Boymaw ORF3 sequence without nonspecific background (Figure 4b). Expression of the Boymaw ORF3 RNA was significantly increased 24 h after starvation (Figure 4c).

**Figure 4.**
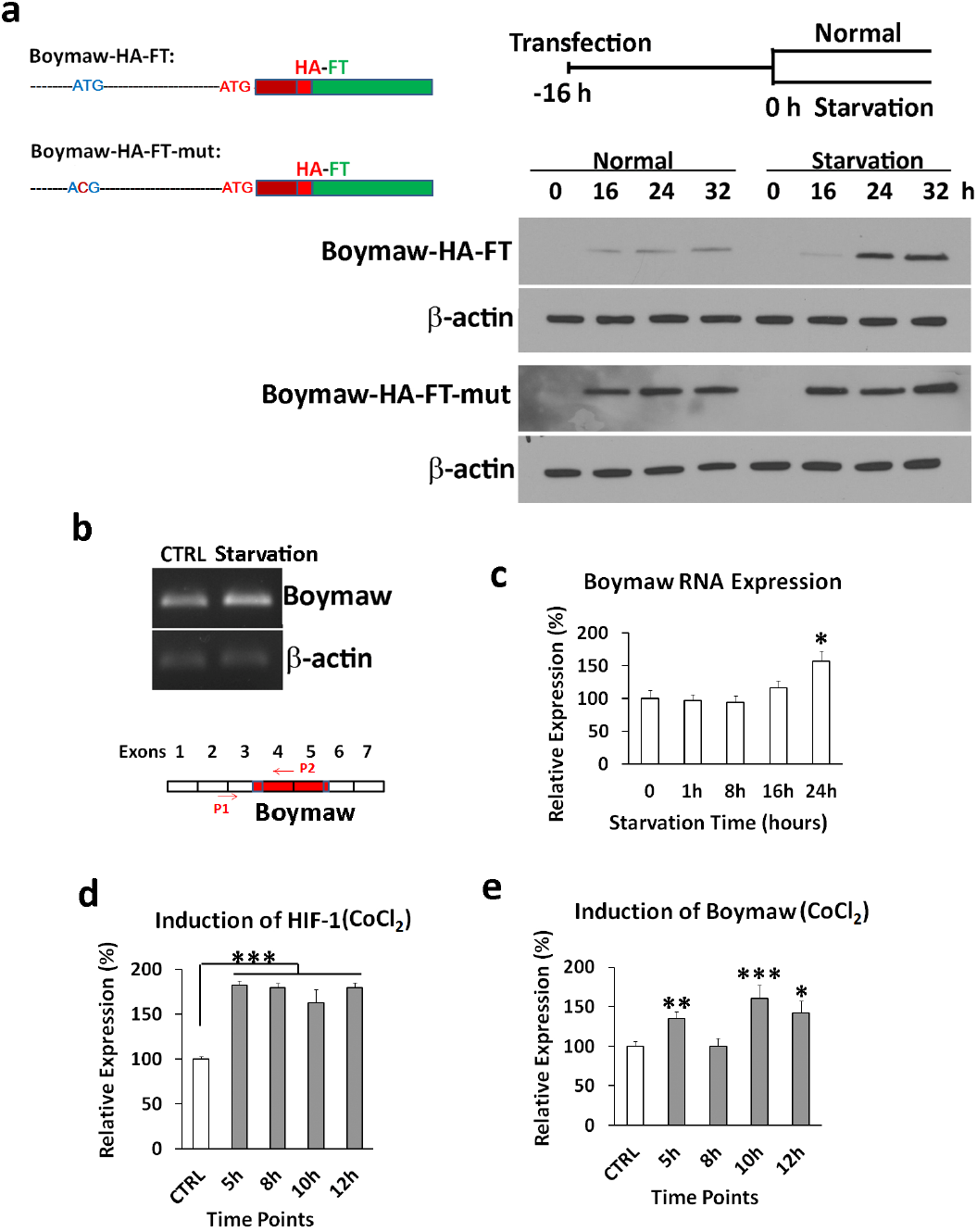
Stress increases Boymaw ORF3 RNA expression. **(a)** Both the Boymaw-HA-FT and the Boymaw-HA-FT-mut are driven by a CMV promoter and terminated by an SV40 poly (A), but differ in a single point mutation of the ATG start codon of the uORF2. Starvation was initiated 16 h after transfection. Expression of the Boymaw-HA-FT fusion protein was examined at 0, 16, 24, and 32 h after starvation. Increase of the Boymaw-HA-FT fusion protein was readily observed at 24 and 32 h post-starvation in cells transfected with the Boymaw-HA-FT plasmid, but not the Boymaw-HA-FT-mut plasmid. **(b)** RT-PCR amplification of the endogenous Boymaw ORF3 RNA in HEK293T cells 24 h after starvation. **(c)** Real-time PCR quantification of the endogenous Boymaw ORF3 RNA expression under starvation. There were 7 to 8 replica wells per treatment per time point. There is a significant treatment effect on the Boymaw ORF3 RNA expression (F(4,34) = 4.79, p < 0.01). *Post hoc* analysis revealed a significant increase of the Boymaw ORF3 RNA expression in cells 24 h after starvation. **(d)** Real-time PCR quantification of the HIF-1α RNA expression at multiple time points after addition of CoCl_2_. A significant effect of treatment was observed (F(4,55) = 28.34, p < 0.00001). *Post hoc* analysis revealed a significant increase of HIF-1α RNA at 5 h, 8 h, 10 h, and 12 h post-treatment. There were 12 replica wells for the treatment per time point. **(e)** In the same set of samples from (d), real-time PCR quantification of the endogenous Boymaw ORF3 RNA expression was conducted. Significant treatment effect was observed (F(4,55) = 6.10, p < 0.001). *Post hoc* analysis revealed significant increase of endogenous Boymaw ORF3 RNA expression in cells treated with CoCl_2_ at 5 h, 10 h, and 12 h post-treatment. There were 12 replica wells for the treatment per time point. Error bar: SEM; * p < 0.05, ** p < 0.01, *** p < 0.001.

Regulation of Boymaw ORF3 translation was also investigated with a different stress, hypoxia. Cobalt chloride (CoCl_2_) was used to induce hypoxia in HEK293T cells [13,14]. The optimal concentration of CoCl_2_ was determined by analysis of induction of hypoxia-inducible factor-1α (Supplemental Figure 3a). The effects of hypoxia on the Boymaw ORF3 translation were investigated in cells transfected with either the Boymaw-HA-FT or the Boymaw-HA-FT-mut construct. Hypoxia had little effect on the induction of the Boymaw ORF3 translation (Supplemental Figure 3b). However, the addition of CoCl_2_ significantly increased RNA expression of hypoxia-inducible factor-1α and endogenous Boymaw ORF3 RNA (Figure 4d and 4e). The significant increase of the Boymaw ORF3 RNA was observed at all different time points except 8 h after addition of CoCl_2_. It is possible that different regulatory mechanisms may be involved in its induction at different times. The exact reason for the induction of two peaks remains to be investigated.

## Boymaw Overexpression in Major Psychiatric Disorders

Expression of the full-length Boymaw ORF3 protein inhibits oxidoreductase activity, rRNA expression, and protein translation. Given that ORF3 protein expression can be induced by stress, we were interested in examining whether expression of the Boymaw ORF3 RNA is increased in the postmortem brains of patients with major psychiatric disorders. The Boymaw ORF3 RNA isoform was quantified, using the same set of the primers described in Figure 4b, in total RNA extracted from human cortex BA7 of Neuropathology Consortium from Stanley Medical Research Institute (SMRI). The demographic information was summarized in Supplemental Table I and at SMRI (http://www.stanleyresearch.org/dnn/Default.aspx?tabid=196). The Boymaw ORF3 RNA expression was quantified against either 18S rRNA or β-actin mRNA control. Significant increase of the Boymaw ORF3 RNA expression was observed in the postmortem brains of patients with schizophrenia regardless of whether 18S rRNA or β-actin mRNA was used as the reference in real-time PCR quantification (Figure 5a and 5b). Increased Boymaw ORF3 RNA expression was not statistically significant in either bipolar disorder or major depression. The relative expression of the Boymaw ORF3 RNA from the two quantification methods was compared at individual patient level (Figure 5c). There was a significant correlation between the relative levels of the Boymaw ORF3 RNA expression in individuals from the two quantification methods. Because both schizophrenia and bipolar patients received antipsychotics (fluphenazine), we investigated whether the increase of Boymaw ORF3 RNA expression was caused by the medication. Linear regression analysis was conducted to investigate the correlation between the amount of fluphenazine taken and the level of the Boymaw ORF3 RNA in the combined schizophrenia and bipolar samples. No correlation was found between the medication and the Boymaw ORF3 RNA expression (Figure 5d). Our studies therefore suggest that increase of the Boymaw ORF3 RNA expression in patients is unlikely caused by the medication. Since age, gender, postmortem brain pH, and postmortem intervals (PMI) were matched between cases and controls in the Stanley collection (Supplemental Table I), it is unlikely that increased Boymaw ORF3 RNA expression in schizophrenia patients was caused by these potential confounding factors. Among many different Boymaw RNA isoforms, expression of the Boymaw ORF3 RNA is very low in healthy human brains. To provide an estimation of its relative abundance, we quantified the level of the Boymaw ORF3 RNA in comparison with the levels of the DISC1 and β–actin mRNAs in human brain tissue. To compare the relative abundance of RNAs between different genes, it is necessary that their real-time PCRs have the same amplification efficiency. We carefully selected two pairs of primers for the DISC1 and Boymaw ORF3 genes that displayed the same PCR amplification efficiency according to amplification of the same molecular concentration of their diluted plasmid templates (Supplemental Figure 4). Human cortex BA7 RNA was pooled from three healthy controls and relative abundance of the Boymaw ORF3, DISC1, and β–actin RNAs was quantified. Our quantification found that DISC1 mRNA is about 40 fold more abundant than the Boymaw ORF3 RNA (Figure 5e).

**Figure 5.**
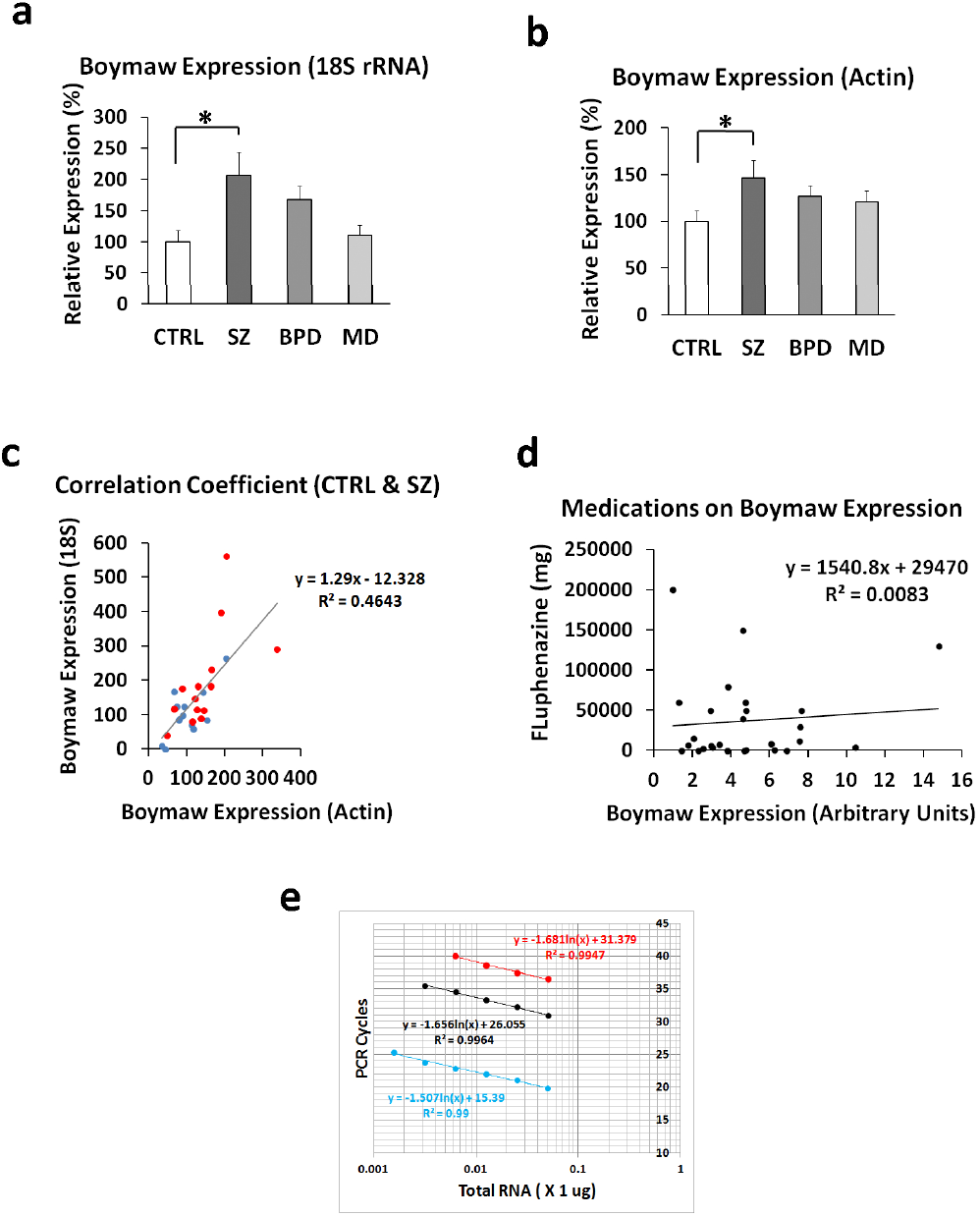
Over-expression of the Boymaw ORF3 RNA in postmortem patient brains. **(a)** Real-time PCR quantification of the Boymaw ORF3 RNA expression in postmortem human brains. The mean value of the Boymaw ORF3 RNA expression from the healthy controls was used to calculate relative expression of the Boymaw ORF3 in schizophrenia (SZ), bipolar disorder (BPD), and major depression (MD). There is a significant diagnostic group effect on the Boymaw ORF3 expression (F(3, 52) = 3.26, p < 0.05). *Post hoc* analysis revealed a significant increase of the Boymaw ORF3 RNA expression in schizophrenia (Tukey test, p < 0.05). Expression of 18S rRNA was used as a reference control to quantify relative expression of the Boymaw ORF3 RNA. **(b)** Expression of the Boymaw ORF3 RNA was quantified from the same samples using β-actin as a reference control to confirm the group difference. A significant increase of the Boymaw ORF3 RNA was observed again (F(3, 51) = 2.71, p = 0.05). *Post hoc* analysis revealed that schizophrenia patients have a significantly increased level of the Boymaw ORF3 RNA expression (Tukey test, p = 0.05). **(c)** Pearson’s correlation analysis revealed a significant correlation between the levels of the Boymaw ORF3 RNA expression quantified relative to 18S rRNA and to β-actin in individual healthy controls and schizophrenia patients. Blue dots: controls; red dots: patients with schizophrenia. **(d)** Antipsychotic fluphenazine has no effect on the expression of the Boymaw ORF3 RNA expression in individual patients (black dots) with schizophrenia and bipolar disorder. **(e)** Quantification of the relative expression of the Boymaw ORF3, DISC1, and β–actin RNA in the healthy human brain. SYBR Green real-time PCR was used to quantify relative expression of the Boymaw ORF3 (red), DISC1 (black), and β–actin RNA (blue) with a comparative Ct method. Error bar: SEM; * p < 0.05.

## Discussion

The Boymaw gene disrupted in the Scottish family has been considered a potential noncoding RNA gene [4–6]. In the present studies, however, we provide evidence to support that the evolutionarily conserved Boymaw ORF3 may encode a small protein. Small functional ORFs are difficult to identify with bioinformatic analysis of the genome. The proteins or peptides encoded by these small ORFs, however, can play important roles in development and disease. For example, the *tarsal-less* gene encodes only 11 amino acids by a short ORF, but the 11-amino-acid-peptide is essential for *Drosophila* development [15]. Our studies found that the Boymaw ORF3 encodes a small protein (63 amino acid residues) which is predominantly localized in mitochondria. Expression of the Boymaw ORF3 protein inhibits MTT reduction, rRNA expression, and protein translation in the same way as the DISC1-Boymaw fusion protein. Interestingly, the Boymaw ORF3 expression is induced by different stressors at RNA and/or protein translation levels. Finally, the Boymaw ORF3 RNA is significantly over-expressed in the postmortem brains of patients with schizophrenia, suggesting that the Boymaw may function as a potential susceptibility gene in the general population of psychiatric patients. Consistent with our hypothesis, duplication of 11q14.3 DNA containing the Boymaw gene co-segregates with major depression in another family [16].

We demonstrated the translation of the Boymaw ORF3 protein using Western blot, immunocytochemical, and iTRAQ analyses. Mutation of the ATG start codon of the ORF3 completely abolished its inhibition in MTT reduction, rRNA expression, and protein translation, further supporting the translation of the Boymaw ORF3. In the human brain, expression of the Boymaw ORF3 RNA is very low and easily outcompeted by other more abundant Boymaw isoforms during PCR amplification [5]. Among other alternatively spliced Boymaw RNAs, it is unknown whether any of them may encode protein. There has been little study on expression of the Boymaw gene in the human brain.

What could be the potential physiological role of the Boymaw ORF3 protein? Under starvation, we found that Boymaw uORF2 is necessary to up-regulate the translation of the downstream ORF3. The exactly same regulatory function by the Boymaw uORF2 was also found in the uORFs of PKCη, a gene involved in cell survival [10]. Under starvation, inhibitory uORFs are required to increase downstream PKCη protein expression. We posit that the Boymaw ORF3 protein induced by starvation is translocated into mitochondria to inhibit oxidoreductase activities to decrease cellular metabolism for cell survival. Starvation also increases expression of the Boymaw ORF3 at the RNA level. However, not all stresses regulate the Boymaw ORF3 expression at both RNA and protein translational levels. Hypoxia significantly increases expression of the Boymaw ORF3 at the RNA level, but has little effect on the translation of the Boymaw ORF3. We speculate that increase of the Boymaw ORF3 protein expression, similar to PKCη protein, could be a part of adaptive responses to stress to promote cell survival (Figure 6). Since persistent over-expression of the Boymaw ORF3 protein may be harmful, rapid degradation of the Boymaw ORF3 protein may be evolved to provide a dynamic regulation of cellular activity and protein translation in respond to changing environments.

**Figure 6.**
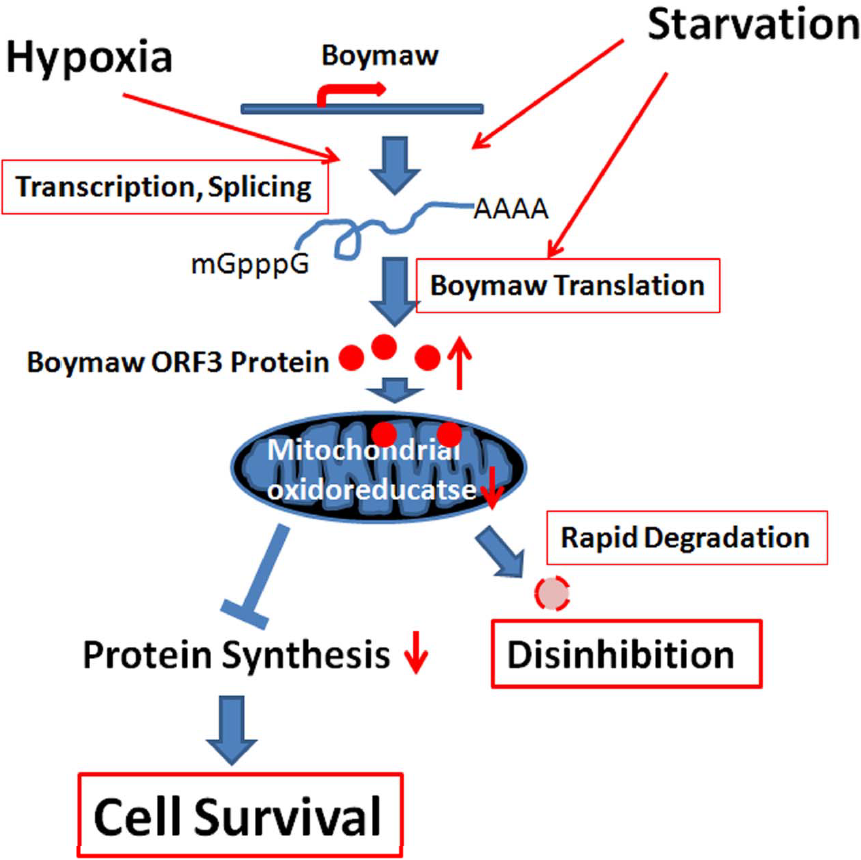
Hypothetical role of the Boymaw ORF3 protein. The Boymaw ORF3 expression can be increased at RNA (transcription and/or splicing) and protein translational levels by different stress. The induced Boymaw ORF3 protein is translocated into mitochondria to inhibit oxidoreductase activity which may further send signals to decrease protein translation to promote cell survival under stress. Rapid degradation of the Boymaw ORF3 protein helps cells to recover quickly in permissive conditions.

Our studies were conducted in HEK293T cells which share 90% of expressed genes with human brain [17,18]. The molecular similarity between the HEK293T cells and human brain may provide support for the relevance of our *in vitro* studies on the functions of the Boymaw ORF3 RNA to the physiological role of the Boymaw gene in human brains. However, our studies have limitations. The Boymaw gene may have different functions in different types of cells; and we do not know which types of cells are the most appropriate for the study of Boymaw functions relevant to the development of psychiatric disorders.

Conceivably, neuronal and glial cells are good candidates, but we cannot rule out the involvement of other types of cells expressing the Boymaw gene. Regardless of cell types, the fundamental biology of the Boymaw gene, that it encodes a small protein which is predominantly localized in mitochondria to inhibit mitochondrial NADH oxidoreductase activity, is unlikely different in different cell types. The consequence of these fundamental functions of the Boymaw gene, however, could vary in different types of cells.

Stress plays an important role in development of psychiatric disorders. Prenatal starvation was reported to significantly increase the risk of developing schizophrenia and other psychiatric disorders [19,20]. Hypoxia was also reported as a risk factor for the development of psychiatric disorders [21]. Environmental stress could partly function through the Boymaw ORF3 pathway to impact brain development and functions via alteration of oxidoreductase activities and protein translation. We found that expression of the Boymaw ORF3 RNA is significantly increased in the postmortem brains of the general population of patients with schizophrenia. However, not all schizophrenia samples display increased expression of the Boymaw ORF3 RNA. There are several possible explanations for this finding. First, genes other than Boymaw are risk factors for patients with normal expression of the Boymaw ORF3 RNA. Second, increased Boymaw ORF3 expression may be transient in patient brains and therefore may not always be detectable in postmortem brains. For example, over-expression of the Boymaw ORF3 RNA can be induced by stress, which may not last very long. Finally, if the Boymaw ORF3 protein is an important labile regulator of cellular metabolic activity, the Boymaw ORF3 RNA may also be short-lived and is susceptible to degradation in postmortem human brains. If over-expression of the Boymaw ORF3 protein is a significant risk factor, a single point mutation of the ATG start codon of the uORF2, which dramatically increases the translation of the Boymaw ORF3 protein, may increase the risk of developing major psychiatric disorders. Rhesus monkeys and baboons naturally carry a mutated (ATC) start codon of the uORF2 in their Boymaw genes. It will be interesting to know whether the uORF2 mutation may present in human psychiatric patients.

Our previous studies reported that expression of the DISC1-Boymaw fusion protein inhibits MTT reduction, rRNA expression, and protein translation (manuscript submitted). Here, we found that expression of the Boymaw ORF3 protein generates the same cellular phenotypes. Since 59 out of 63 amino acid residues of the Boymaw ORF3 are in-frame fused to the C-terminus of the truncated DISC1 protein in the Scottish family, the Boymaw ORF3 protein is therefore a potential mechanistic link between the DISC1-Boymaw fusion protein in the Scottish family and the Boymaw ORF3 over-expression in the general population of patients with schizophrenia. In the Scottish family, the fused Boymaw ORF3 protein is likely over-expressed in the form of the DISC1-Boymaw fusion proteins. Two lines of evidence support this hypothesis. First, we found that endogenous DISC1 mRNA is about 40 fold more abundant than the Boymaw ORF3 RNA in healthy human brain. In the Scottish family, the DISC1-Boymaw fusion RNA, driven by the endogenous DISC1 gene promoter, will be likely more abundant than the endogenous Boymaw ORF3 RNA, even though the DISC1-Boymaw fusion transcript is 2 to 3 fold less than wildtype DISC1 mRNA [6]. Second, the DISC1-Boymaw fusion RNA contains a single ORF optimal for efficient protein translation while the endogenous Boymaw ORF3 translation is inhibited by its overlapping uORF2. The translocation, however, abolished both inhibitory mechanisms of the Boymaw ORF3 expression at RNA and protein translational levels by the formation the DISC1-Boymaw fusion gene. Therefore, we propose that over-expression of the Boymaw ORF3 protein, through different pathways, could be a potential common mechanism between the Scottish family and the general population of patients to inhibit protein translation and thereby contribute to the pathogenesis of major psychiatric disorders.

## Acknowledgment

We thank Stanley Medical Research Institute to provide RNA samples from the postmortem brains of patients with psychiatric disorders.

### Contributions

X.Z. conceived and wrote the studies. B.J., M. K., K.H. performed and analyzed the research data.

